# High Temperature and Microbiome Conditions Affect Gene Expression in Soybean

**DOI:** 10.1101/2024.11.04.620947

**Authors:** Liza Van der Laan, Dinakaran Elango, Antonella Ferela, Jamie A. O’Rourke, Asheesh K. Singh

## Abstract

Heat stress is increasingly a problem in global agriculture production, both in increasing occurrences and extended durations. Understanding the molecular mechanisms of the soybean heat stress response is essential for breeding heat tolerant soybeans. Plant associated microbiomes are known to mitigate adverse effects from abiotic stress. Soybean heat stress studies have primarily focused on response to short periods of stress, and how soybean responds on a transcriptional level to a soil microbiome is poorly understood. We hypothesize a soil microbiome may help soybean survive long-term heat stress exposure. We used RNA-seq to measure the transcriptional responses in four soybean exposed to two temperature regimes and grown in two soil microbiome conditions. We identified unique responses to temperature based on the soil microbiome conditions and to the different genotypes, with fewer changes across genotypes in response to a soil microbiome. Our findings provide insights on the interaction of soil microbiome with heat stress response in soybean and identify gene targets to further study the soybean heat stress tolerance with applications to develop improved varieties.

## Introduction

Agriculture production is vulnerable to abiotic stress, which is becoming more prevalent due to climate change. Extreme weather events are becoming increasingly common, with the long-term forecast of climate conditions showing an increase in the number and duration of heat waves (Perkins-Kirkpatrick and Lewis, 2020), as well as an increase in average air temperatures (IPCC, 2023). These conditions threaten many crops around the world, including soybean [*Glycine max* L. (Merrill)], the worlds largest oilseed crop, valued for high protein and oil content(Cober et al., 2009). Few mitigation strategies exist to cope with heat stress; therefore, the development of heat-tolerant crop cultivars is essential. Understanding of the mechanisms associated with heat stress responses behind can aid and accelerate breeding efforts.

Heat stress negatively affects crop production by reducing plant growth and lowering seed production, resulting in yield loss (Cohen et al., 2021; Deryng et al., 2014). Plants may be tolerant or susceptible to heat stress, with susceptible plants more likely to experience cellular damage or localized cell death, which can lead to a metabolic imbalance (Bita and Gerats, 2013; Mittler et al., 2012). Plants have developed a variety of adaptations to mitigate the effects of heat stress and protect from further damage, including activating a heat stress response network (Mittler et al., 2012). Adaptations to heat stress include observable changes in the leaf orientation and early maturation, while other responses include modification of membrane compositions, accumulation of osmoprotectants, and an increased synthesis of secondary metabolites (Bita and Gerats, 2013). Control of these responses is finely regulated at all stages of gene expression, from transcription to post-translational modifications (Guerra et al., 2015).

Plants have three main microbiomes: the soil microbiome, the above-ground microbiome, and the intercellular microbiome. Microbiomes are dynamic, with their structures and functions changing due to changes in the surrounding environmental conditions (Timm et al., 2018). All three are hypothesized to promote plant growth and to assist plants tolerance of different sources of stress (Rivero et al., 2022), although the soil microbiome has been the most studied. Soil microbiome plays a key role in plant development, reproduction, and health (Berendsen et al., 2012; Fitzpatrick et al., 2018). Changes in the soil microbiome are also driven by an interaction with the surrounding plants, which seem to actively seek cooperation and increases in specific types of microorganisms during times of abiotic and biotic stress (Bakker et al., 2018). These interactions between the plants and soil microbiome are thought to increase the plant tolerance to different abiotic and biotic stresses, including heat stress (Fitzpatrick et al., 2018; Stringlis et al., 2018). Limited knowledge exists on the interaction between plants and microbiomes and how microbiomes may assist plants during times of stress. Information on how microbes interact with the plant, including changes in gene expression and what microbial species are the most prevalent, would be useful in ensuring a microbiome that promotes stress resilience in plants.

RNA-sequencing (RNA-seq) is a powerful tool to study plant abiotic stress responses. In soybean, gene expression studies have investigated the response to multiple stresses including drought (Chen et al., 2016; Rodrigues et al., 2015), flooding (Chen et al., 2016), elevated ozone (Whaley et al., 2015), nutrient deficiency (Kohlhase et al., 2021; Moran Lauter et al., 2014; Zeng et al., 2018; H. Zeng et al., 2019), salinity (Cadavid et al., 2020; A. Zeng et al., 2019), and cold (Hussain et al., 2023). Additionally, there are studies examining heat stress in soybean with three studies focusing on gene expression of soybean undergoing short-term heat stress treatments in both the leaf tissue (Song et al., 2017; Wang et al., 2018) and in the roots (Valdés-López et al., 2016). Song et al. (2017) studied two different soybean genotypes undergoing short-term heat stress and found that the genotypes differed in their gene expression patterns, including 51 genes identified with inverse regulation between the two genotypes.

They further focused on the expression of heat shock factors (HSFs) and found these to be highly expressed under short periods of heat stress and then decreased in their expression past 12 hours (Song et al., 2017). In another study examining leaves in a single genotype, gene ontology (GO) term enrichment of DEGs was identified in categories related to transport, response to stress, response to stimulus, and binding (Wang et al., 2018). This study found that 11 TF families accounted for the majority of the stress-response TF genes, including the bHLH, MYB, HSF, NAC, WRKY, and bZIP families (Wang et al., 2018). One study looked at transcriptional responses for longer term heat stress while in reproductive growth stages and found an overexpression of HSF and unfolded protein response pathways in response to heat, with HSFA2 mainly being expressed in reproductive tissues (Sinha et al., 2023). A proteomics study analyzed mechanisms for soybean in heat, drought, and combined stress and found that many of the differentially expressed proteins were associated with photosynthesis and ATP synthesis (Das et al., 2016). The transcriptome profiling of soybean under heat stress has focused on short exposures, with most studies only exposing for up to 24 hours and the longest exposure being 15 days. Studies in *Arabidopsis* have shown that the cellular responses and gene expression levels are different between short-term and long-term heat stress (Tang et al., 2016). Given that climate change is likely to increase the average air temperature, it is essential also to understand how soybean responds to long exposures to heat stress as well as shorter exposures.

The objective of this study was to compare differential gene expression patterns of multiple genotypes of soybean under a long period of continuous heat stress. Additionally, this is one of the first reports to investigate the transcriptional interaction of multiple soybean genotypes in abiotic stress conditions with the soil microbiome. We used RNA sequencing on leaf tissue from four genotypes grown continuously in either control or heat stress conditions in both the presence and absence of a native soil microbiome. These results indicate a difference in genotype response to heat stress and that the soil microbiome interacts with this response. This study provides insight into heat response genes, as well as the different responses across genotypes and the impact of the microbiome on the response.

## 2. Materials & Methods

### 2.1. Soil Preparation

A mixture of field soil and sand was used for sowing. The field soil was sourced from the Burkey Research Farm from Iowa State University (ISU) [42°0’50”N, 93°47’30”W] and is a Clarion loam. Topsoil from the field was collected and run through a pulverizer to ensure an even soil texture. The soil was sieved to remove any stones and field debris. Half of the soil was then autoclaved to eliminate the microbiome of the soil. Soil samples were taken for further analysis of soil nutrients, as well as quantification of microbes in the native and autoclaved soils. Using a small concrete mixer, equal amounts by volume of field soil and construction sand (ISU Facilities Planning and Management) were mixed to get the soil mixture. Mixtures that included the autoclaved soil were autoclaved after mixing to eliminate any microbes introduced due to the construction sand or the concrete mixer.

### 2.2. Plant Materials & Treatments

Four genotypes of soybean were utilized in this experiment: (1) Williams82 (PI518671), (2) IAS 19C3, (3) PI89008, and (4) PI639693. Williams82 (MG III) was selected as the control genotype due to the availability of its reference genome. IAS19C3 (MG II) was selected as an elite variety out of the ISU soybean breeding program. The two PI lines were selected as heat tolerant (PI639693 [MG II]) and as heat susceptible (PI89008 [MG II]) based on prior screening. Seeds from the four genotypes were surface sterilized and sown into pots filled with the soil mixture outlined above. Pots were immediately placed into assigned growth chambers at the ISU Enviratron facility (Bao et al., 2019). Chambers were programmed at treatment temperatures the day before sowing to allow for proper temperatures at the time of sowing. The temperatures were 28/21 (d/n) for the control and 38/28 for the heat stress, and 60% relative humidity and 400 ppm CO2. Lights were on a 16/8 hour day and night schedule. Two chambers were set at each temperature, and one soil type was placed in each chamber for a total of a four-factorial treatment design. Four seeds were sown per pot and thinned to one when the unifoliate leaves were unfolded (VC stage). Four biological replicates of each genotype in each temperature-soil combination were planted in individual pots.

All pots were watered daily to ensure no drought conditions were introduced. Ultra-pure water from the Thermo Scientific GenPure Pro system was used to ensure that no microbes were introduced to the soil microbiome via watering. At 33 days after sowing, growth stages (Fehr et al., 1971) were recorded. The emerging trifoliate, including the shoot apical meristem, was collected, flash-frozen in liquid nitrogen, and then maintained at -80 °C. This tissue is herein referred to as the leaf tissue. All four biological replicates of each genotype and treatment were sampled.

### 2.3.RNA Extraction & Sequencing

Total RNA was extracted and isolated from the leaf tissue using the RNeasy Plant mini kit (Qiagen, Cat#74904), following manufacturer’s protocols. The RNA quantity and quality were checked with a Qubit 2.0 Fluorometer (Thermo Fisher Scientific, Waltham, MA). RNA samples were sent to the

Iowa State University DNA Facility for library preparation and sequencing. Library construction was with QuantSeq 3’mRNA, and sequences were generated on the NovaSeq 6000 (Illumina, San Diego, CA) with 100 bp single end reads.

### 2.4. Identification of Differentially Expressed Genes

Sequence data quality was checked using FastQC (Andrews, 2016). Trimmomatic was used to remove reads with a quality score lower than 20 and sequencing adaptors. Cleaned fastq files were mapped to the soybean reference genome (*Glycine max* Wm.82.a4.v1, Phytozome) using STAR (version 2.7.1 (Dobin et al., 2013)) using the Iowa State University Nova high-performance computing cluster, identifying 37,838 expressed genes. SAMtools (version 1.16.1 (Danecek et al., 2021)) was used to sort the mapped reads, and the resulting binary alignment/map (BAM) files were used for further downstream differential expression analyses.

Differential expression analysis was performed in RStudio. The edgeR package (Robinson et al., 2009) was used to identify differentially expressed genes (DEGs). Library sizes were normalized across all samples within tissue type using the trimmed mean of M-values (TMM) method (Robinson and Oshlack, 2010). A design matrix was built using the normalized count data with genotype, soil type, and temperature treatment as factors. Simple comparisons were made as contrasts between a single genotype in a contrasting environment (either changing temperature or changing soil microbiome).

Complex comparisons via individual contrast statements were made between temperature conditions and genotypes, as well as between soil microbiome and genotypes. The likelihood ratio test was used for each contrast to test for differential expression of the temperature or soil microbiome effect with each genotype compared to Williams82. Contrast statements for four types of complex analyses include

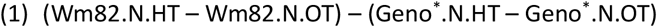

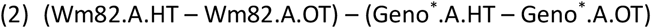

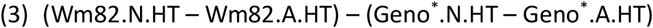

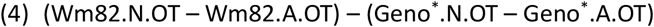

Where N is non-autoclaved soil with a native soil microbiome, A is autoclaved soil with no microbiome, OT is optimal temperature, HT is heat stress temperature, and Geno represents the three genotypes that were compared to Williams82. Genes with a false discovery rate (FDR) <0.05 were considered differentially expressed.

### 2.5. Gene Annotation, Gene Ontology, & Transcription Factors

Gene annotation for the DEGs were annotated using the Gene Annotation Lookup tool at SoyBase.org. Overrepresented (corrected p-value <0.05) gene ontology (GO) terms related to biological processes were identified from the DEG lists using the GO Term Enrichment Tool at SoyBase.org (Morales et al., 2013). Transcription factors (TF) were identified using the PlantTFDB database (https://planttfdb.gao-lab.org (Jin et al., 2016)).

### 2.6. Gene Co-expression Network Analysis

To construct weighted co-expression networks, we used the R package CEMiTool (version 1.24.0 (Russo et al., 2018)). Two networks were built to explore the gene co-expression patterns to heat stress in different soil microbiome conditions: (1) heat stress in native microbial soil and (2) heat stress in autoclaved soil. For all networks, only DEGs that were significant for all genotypes were used to construct the networks. Because of the low number of DEGs across all genotypes for the effect of soil microbiome analyses, we elected not to construct networks for these comparisons. A normalized count table from all these genes lists was used as input. Over representation analysis was conducted using GO biological process terms downloaded from SoyBase.org for each gene list. Gene interactions in each co-expression module were visualized in CEMiTool using the full integrated soybean network data available on SoyNet (https://www.inetbio.org/soynet/ (Kim et al., 2017)).

## 3. Results

### 3.1. Physiological changes of soybean due to heat stress and soil microbiome

We studied the physiological response of soybean under heat stress and with differences in the soil microbiome. The temperature treatment did not affect the growth stages when the plants were grown in native soil. However, the temperature treatment did have a significant effect (p < 0.05) when the plants were grown in the autoclaved soil. IAS19C3 and PI89008 had reached a significantly later growth stage in the heat treatment (V6) as compared to the optimal treatment in which these genotypes were at V4 and V5, respectively. The soil treatment affected the growth stage as well. When grown in heat, the plants sown in autoclaved soil had earlier growth stages, with Williams82 and PI639693 having significantly earlier stages (V5) as compared to the native soil in which these genotypes were at V6 and V7 stages, respectively. The optimal temperature treatment resulted in all four genotypes having earlier growth stages when grown in autoclaved soil compared to the native microbial soil, with growth stages in the autoclaved soil being two V stages behind the native soil.

### 3.1. Effect of Temperature

#### 3.1.1. Identification and Comparison of DEGs across Genotypes

We studied the effect of heat stress by comparing the expression of genes from plants grown under heat stress compared to plants grown under optimal (control) conditions for each genotype. Due to the multiple factors in this experiment, we performed these analyses separately for each soil type (autoclaved(_a). native microbes (_m)), as well as an additional analysis where soil microbiome type (autoclaved or native microbes) was ignored. The number of DEGs (FDR < 0.05) ranged from 0 to 9128 DEGs, depending on the genotype (Figure 2). The genotypes IAS19C3 and PI639693 had the lowest number of DEGs in response to heat stress, while Williams82 and PI89008 had the highest number of DEGs. DEGs in Williams82 were detected only in native microbial soil and the combined soil analysis, while for PI89008, the DEGs were primarily identified in the autoclaved soil analysis. When comparing the DEGs between each genotype, there was minimal overlap (Figure 3). Only the combined soil analysis identified DEGs shared between genotypes, with the greatest overlap of 112 DEGs being between

**Figure 1.**
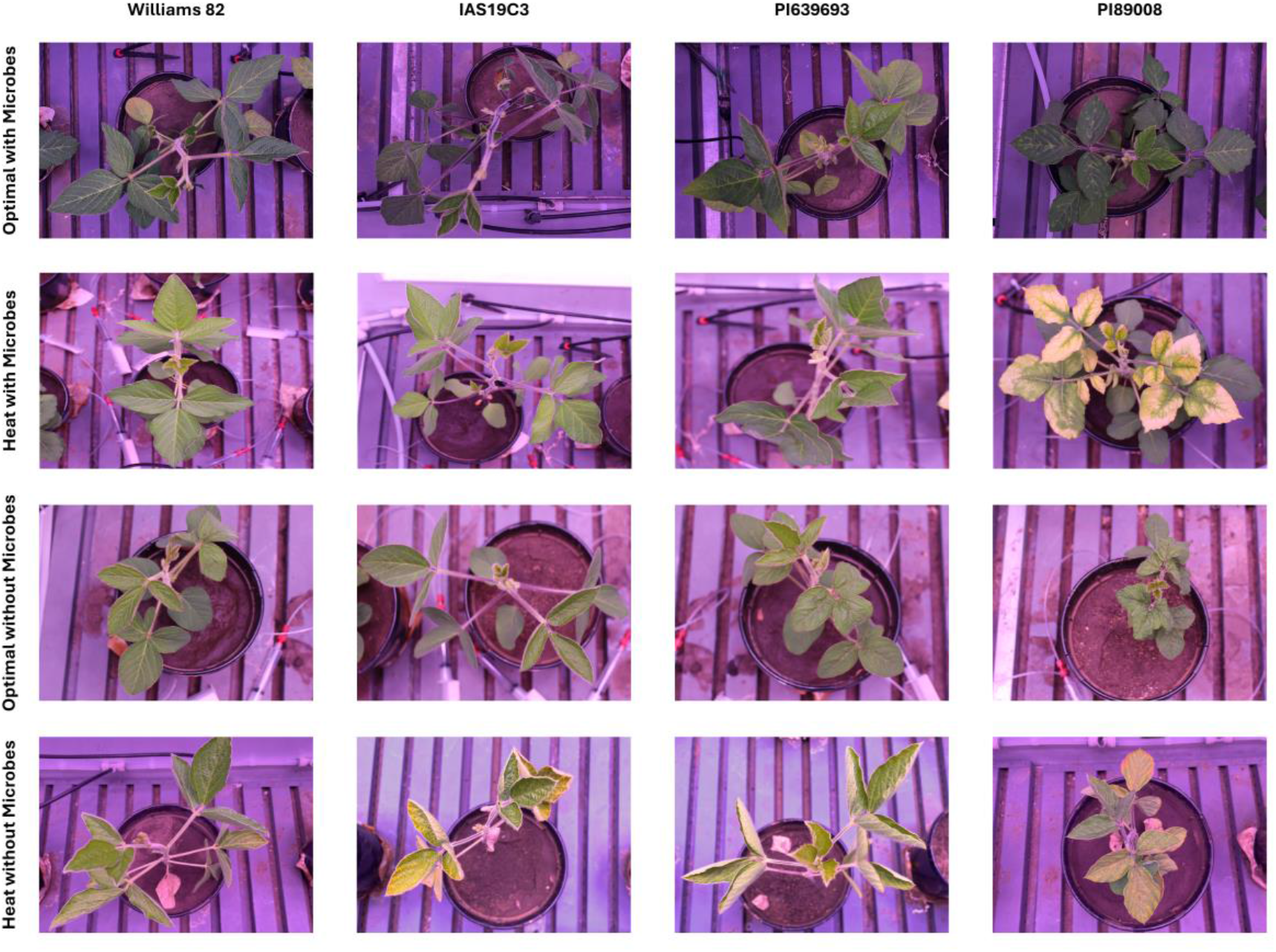
Images of a representative plant from each genotype grown in each temperature-soil microbiome combination. Williams82 is a reference genotype, IAS19C3 is an elite genotype, PI639693 is a heat tolerant genotype, and PI89008 is a heat susceptible genotype. All images were taken 33 days after planting on the day of tissue sampling.

**Figure 2.**
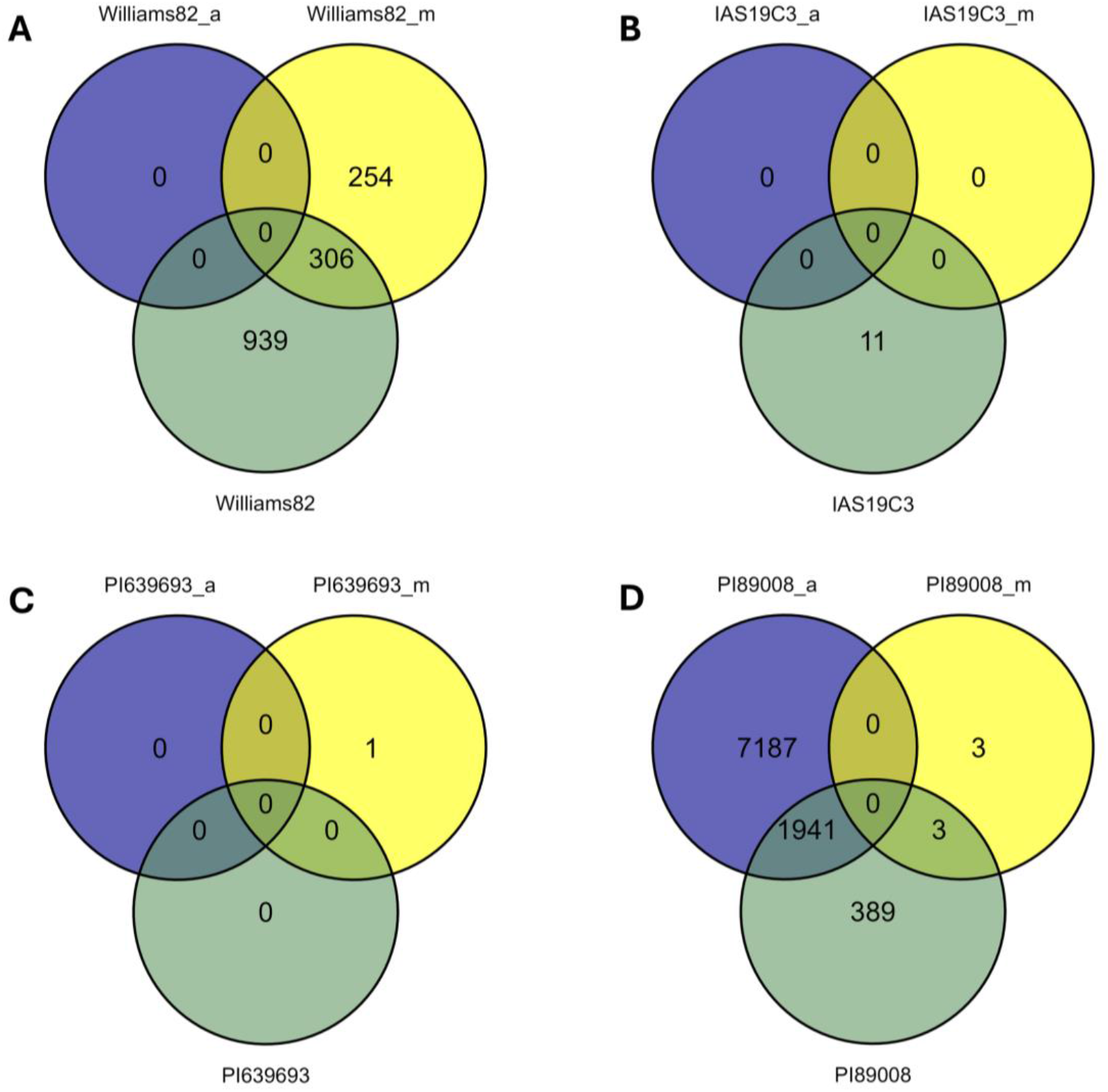
Venn diagrams detailing the distribution of significant DEGs (FDR < 0.05) due to the temperature treatment (heat stress vs control) for each genotype. Each genotype compares the DEGs identified when grown in autoclaved soil (_a), when grown in native microbial soil (_m), and the analysis of combined soil types.

**Figure 3.**
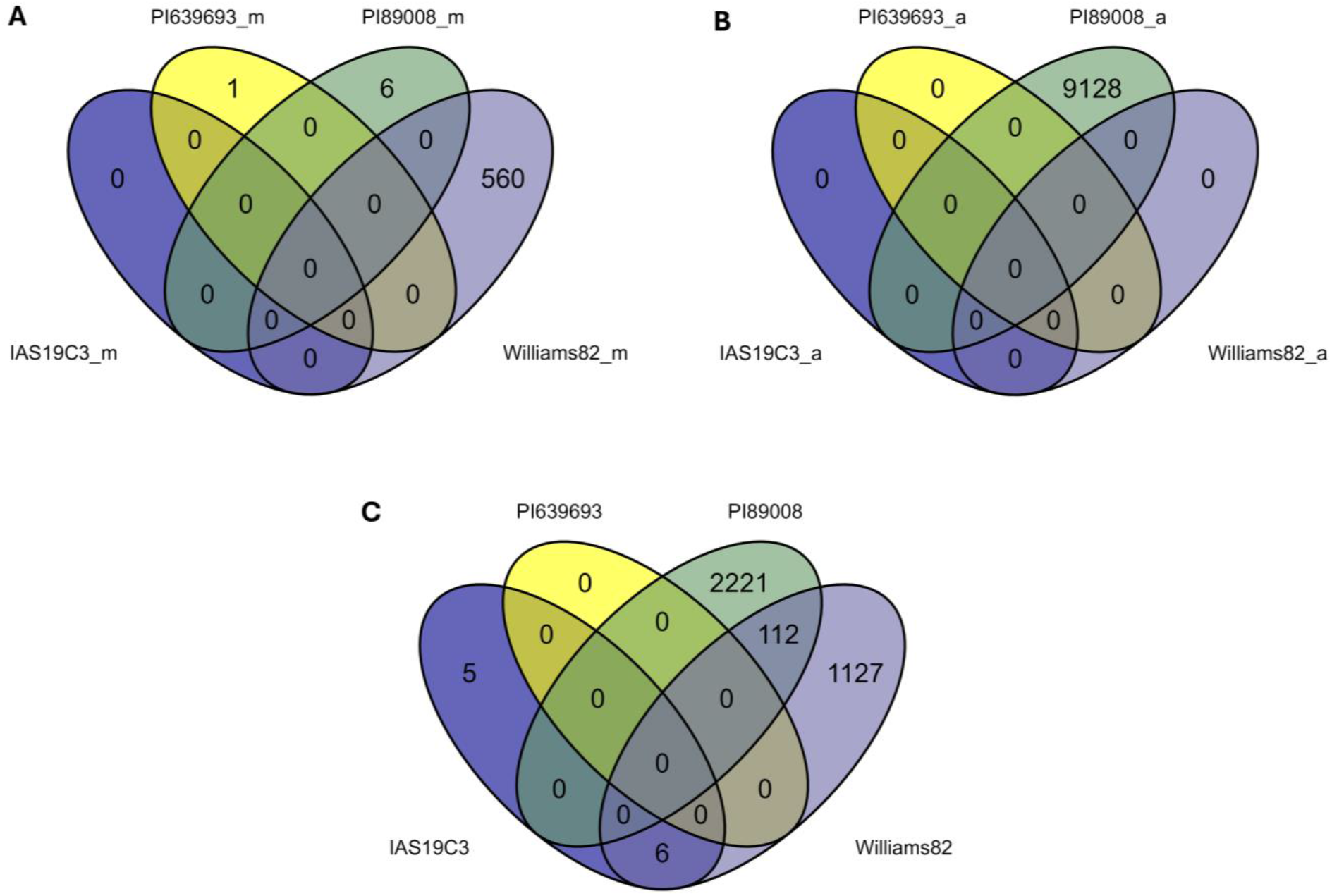
Venn diagrams of the distribution of overlapping DEGs due to temperature treatment for each genotype. Here, we show the overlap between genotypes when using the same soil treatment, with (A) being the native microbial soil, (B) being the autoclaved soil, and (C) being the combined analysis across soil types.

Williams82 and PI89008 and then another six DEGs between Williams82 and IAS19C3. These results suggest differences in heat stress responses across genotypes.

Of the six overlapping DEGs between IAS19C3 and Williams82 (Figure 3), *Glyma.06G289600* was upregulated in both genotypes. This gene encodes for a peptidase family M48 protein and is homologous with the overlapping activity with the *m*-AAA protease 1 (OMA1) gene in *Arabidopsis*. This gene has been found to increase the tolerance of plants to long-term moderate heat stress via reduction of protein aggregation in the mitochondria (Maziak et al., 2021). The remaining five overlapping genes were all downregulated in both genotypes. Several of these genes have been found to be involved in various abiotic stress responses in plants (Ambroise et al., 2024; Castañeda et al., 2021).

The 112 overlapping genes between Williams82 and PI89008 (Figure 3) include 16 upregulated genes that encode for heat shock proteins (HSPs), with the majority belonging to the HSP20 family. Two tandem HSP genes, *Glyma.14g099900* and *Glyma.14g100000*, are homologous to *HSP17.6II*, which is involved in the miR160 pathway regulating heat stress responses in *Arabidopsis* (Lin et al., 2018). Two other upregulated genes, *Glyma.09g131500* and *Glyma.16g178800*, encode for *GmHsp90A1* and *GmHsp90A2*, respectively. Overexpression of *GmHsp90A2* has been found to reduce the degradation of chlorophyll and reduce lipid peroxidation in soybean undergoing heat stress (Huang et al., 2019). This same study found that *GmHsp90A2* interacts with *GmHsp90A1* and that *GmHsp90A1* may play a role in the entry of *GmHsp90A2* into the nucleus (Huang et al., 2019). In a previous study, these four genes were all found to be upregulated in exposure to eight hours of heat stress, and three of these genes were found to still be upregulated after 24 hours of heat stress (Wang et al., 2018). An additional upregulated gene, *Glyma.17g227600*, is homologous with *HSFA2* in *Arabidopsis* which plays a major role in heat stress tolerance by promoting synthesis of jasmonic acid (JA) (Guo et al., 2024). GO term enrichment of these 112 shared genes resulted in a significant overrepresentation of five GO biological process terms: response to heat (GO:0009408), response to high light intensity (GO:0009644), response to hydrogen peroxide (GO:0042542), protein folding (GO:0006457), and response to endoplasmic reticulum stress (GO:0034976). Heat shock factors were identified in each of the over-represented GO terms, with the majority (7 HSPs) represented in the response to heat GO term.

#### 3.1.2. Identification and Comparison of DEGs between Genotypes

Because of the indication of genotypic differences, we wanted to compare different genotypes grown in the contrasting temperature treatments. For these comparisons, we considered Williams82 the control genotype due to the availability of its reference genome. The remaining three genotypes were compared to Williams82 (Equations 1 & 2). These analyses identified a greater number of DEGs than the simple single genotype comparisons, with all three genotype-temperature comparisons identifying DEGs. In both soil types, PI639693 had the fewest number of DEGs detected, indicating that it was the most similar to Williams82 in its response to heat. IAS19C3 had the highest number of DEGs for response to heat when grown in the native microbial soil, while PI89008 had the highest when grown in the autoclaved soil. There was an overlap of 4224 and 964 DEGs (Figure 4A & B) between the three genotype-temperature comparisons when grown in native and autoclaved soil, respectively. All overlapping genes were regulated in the same direction across the three genotypes. From the 4224 and 964 overlapping DEGs between all three genotypes, 83 were shared between the autoclaved analyses (Figure 4B) and the microbiome analyses (Figure 4A). When comparing the DEGs from single genotypes across the soil microbiome conditions, we found 605, 200, and 1226 shared genes for IAS19C3, PI639693, and PI89008, respectively. The DEGs were not all regulated in the same direction, and different regulation trends were noted based on genotype. The majority of shared DEGs across soil microbiome conditions for IAS19C3 and PI639693 were regulated in the same direction, while for PI89008, the majority of the shared DEGs were regulated in opposite directions.

**Figure 4.**
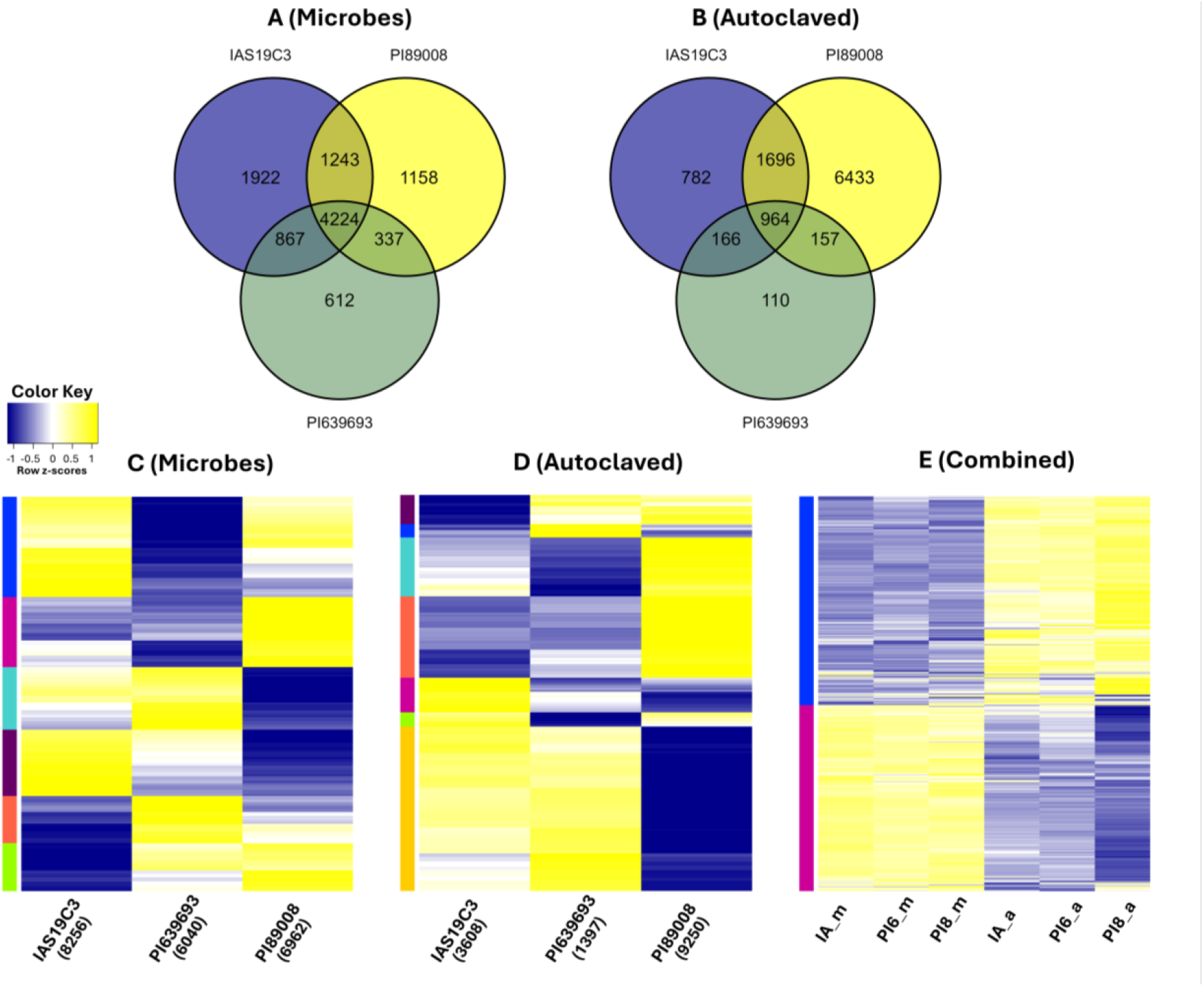
Distribution of differentially expressed genes (DEGs) and heatmaps of the DEGs. The Venn diagrams are the distribution of statistically significant (FDR < 0.05) DEGs in each of the analyses, with (A) being the effect of temperature and genotype when plants were grown in native microbial soil and (B) being when the plants were grown in autoclaved soil. Genotypes were compared to the control of Williams82 in optimal temperature conditions. The heatmaps demonstrate the expression levels of DEGs across all genotypes and analyses, with (C) being the DEGs of the effect of temperature and genotypes with plants grown in native microbial soil, (D) when the plants were grown in autoclaved soil, and (E) the DEGs across the microbial and autoclaved soil analyses. Expression of all DEGs identified for each analysis is represented, even if a gene was not statistically differentially expressed in a specific comparison. The number of significant DEGs for a single genotype is in parentheses below the genotype name. Downregulated genes (negative Z-scores) are shown in blue, and upregulated genes are shown in yellow. Clusters of genes with similar expression profiles are outlined in colors to the left of each heatmap.

We proceeded to primarily focus on the 964 and 4224 DEGs that were significant for all three genotypes in autoclaved and microbially intact soils, respectively. The 964 DEGs were identified from plants grown in autoclaved soils. GO enrichment analysis identified ten statistically significantly overrepresented GO terms among the 964 DEGs, eight of which are associated with photosynthetic processes. The remaining two GO categories are response to heat (GO:0009408) and protein folding (GO:0006457). In contrast, the analyses identified 4224 DEGs common to all three genotypes in non-autoclaved soils. GO analysis of the 4224 genes identified eight over-represented GO terms, including protein folding (GO:0006457) and GO:0009773, which is associated with photosynthetic electron transport. The remaining GO terms are response to heat (GO:0009408), response to hypoxia (GO:0071456), response to salicylic acid (GO:0009751), response to abscisic acid (GO:0009737), regulation of meristem growth (GO:0010075), and defense response to bacterium (GO:0042742). The prevalence of photosynthetic genes identified from leaves of plants grown in autoclaved soils compared to the diverse GO categories identified in leaves of plants grown in microbially robust soils indicates the

microbes actively participate in influencing plant gene expression patterns under heat stress conditions. Given these findings, we were interested in examining the distribution of genes encoding HSPs. Our analyses identified 17 HSPs in leaves from the autoclaved soil and 49 in leaves from the microbial soil. The majority of the HSPs were upregulated due to heat stress, although there were one and five HSPs downregulated in the autoclaved and microbial treatments, respectively. The majority of the HSPs were HSP20s, followed then by HSP70s, and finally HSP90s. The four HSPs that were found in the within genotype comparisons (*Glyma.14G099900, Glyma.14G100000, Glyma.09G131500*, and *Glyma.16G178800*) were also differentially expressed in the between genotype comparisons. Notably, *Glyma.09G131500* was upregulated across all genotypes in all soil types, while the remaining three genes were all upregulated but not in all comparisons. The HSF *Glyma.17G227600*, which was found in both the within genotype and between genotype comparisons, was upregulated across all genotypes and soil types.

Comparing the DEGs identified in all three genotypes grown in autoclaved and microbial soils identified 83 DEGs that were shared across all three genotypes from both soil sources. Examining the expression of these genes found that 62 were expressed in the same direction from both soil sources, while 21 were expressed in the opposite direction, depending on soil sources. GO analyses identified two overrepresented GO terms: response to heat (GO:0009408) and protein folding (GO:0006457).

Several genes of interest identified in previous soybean heat stress tolerance GWAS (Van der Laan et al., 2024) were identified in at least one analysis. Two downregulated DEGs from PI89008 are within 50 kbp from a marker detected for a soybean shoot biomass heat stress index. These DEGs, *Glyma.10G219100* and *Glyma.10G219200*, are homologs for laccase proteins, which are involved in plant growth and development. An additional upregulated DEG, *Glyma.10G217900*, is further away from the previously reported marker. This gene was previously reported to be upregulated when exposed to 24 hours of heat stress (Wang et al., 2018) and is homologous with the *Arabidopsis* gene *AtSUC2*, which is a sucrose carrier that is involved in sucrose loading and found to be essential in plant development (Stadler and Sauer, 2019). Another gene, *Glyma.04G242700*, was upregulated in PI89008 when grown in heat conditions with autoclaved soil and downregulated in microbial soil. This gene was previously found to be associated with shoot and root biomass in heat stress (Van der Laan et al., 2024). This gene is homologous with *AtSKP2B*, which is an F-box protein found to be involved in cell division, which is

important for the development of leaves, meristems, and lateral roots (del Pozo et al., 2006; Manzano et al., 2012).

To visualize the different responses of the genotypes to temperature and their significant DEGs, we generated heatmaps (Figure 4C-E). In total, there were 10,363 and 10,308 significant DEGs across the three genotypes when grown in native microbial soil and autoclaved soil, respectively. The heatmap for the genotype-temperature comparison in microbial soil was organized into six expression clusters. In contrast, the heatmap for the comparison in autoclaved soil was organized into seven expression clusters. GO term enrichment analysis (Corrected P-value < 0.05) was used to determine if the expression clusters were linked with specific biological functions. Of the six clusters from the comparisons grown in the microbial soil, all clusters had one to 32 enriched GO terms. Three of the seven gene clusters for the plants grown in autoclaved soil had no enriched GO terms. The remaining four clusters ranged from one to 58 enriched GO terms. When we generated a heatmap comparing the DEGs across both soil microbiomes (Figure 4E), two clear clusters were identified. These clusters had 37 and 74 significant GO terms, respectively, with a single GO term significant in both clusters (GO:0009408; response to heat).

#### 3.1.3. Differentially Expressed Transcription Factors

Using PlantTFDB, we identified transcription factors (TFs) among the DEGs as genes of interest for further analysis. The log fold-change of the differentially expressed TFs grouped by transcription factor family (TFF) by genotypes and soil treatment is shown in Figure 5. In the autoclaved soil, we identified 650 unique differentially expressed TFs, belonging to 50 TFFs. Most of these differentially expressed TFs were unique to a single genotype, with 74% being differentially expressed in a single genotype, 16.2% differentially expressed in two genotypes, and only 9.8% differentially expressed in all three genotypes. The majority of these TFs (60%) were upregulated due to heat stress. Plants grown in soil with microbes had a greater number of differentially expressed TFs, with a total of 944 TFs from 52 different TFFs. In contrast to the TFs in the autoclaved soil, the majority of the differentially expressed TFs in the microbial soil were shared between the three genotypes (48.8%), with 20.9% shared between two genotypes and 30.3% being unique to a single genotype. The majority of the TFs (73%) in the microbial soil analysis were downregulated due to heat stress.

**Figure 5.**
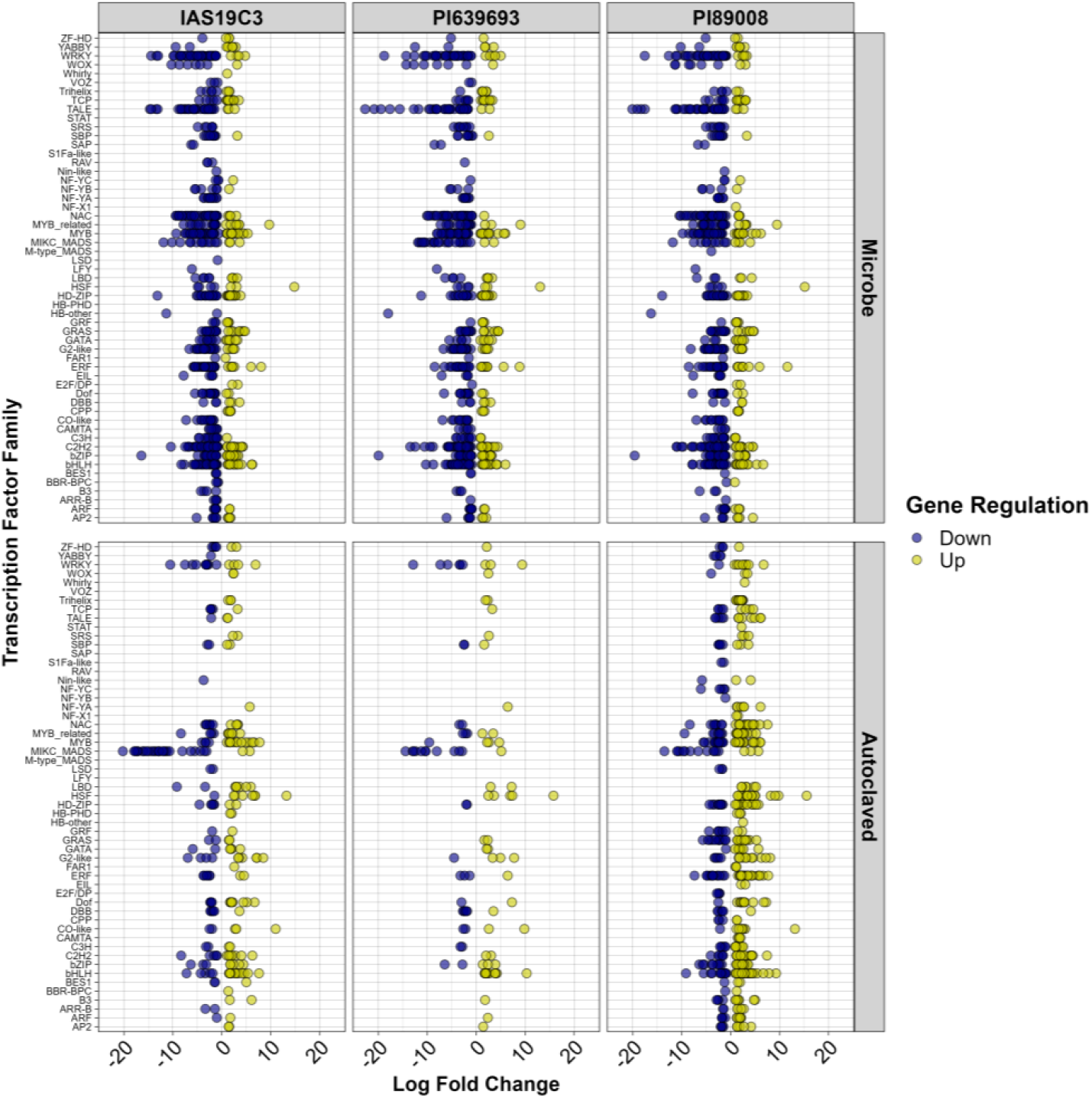
Expression values of differentially expressed transcription factors of three soybean genotypes compared to the control genotype Williams82. Differentially expressed genes were identified in the leaf tissue of plants in response to prolonged heat stress. Plants grown in heat were also grown in two different soil microbiome conditions: native microbial and autoclaved soil. Transcription factors were identified in each DEG list and plotted by transcription factor family using the log fold-change of the gene. Up-regulated genes are shown in yellow, while down-regulated genes are shown in blue.

Between the two soil microbiome analyses when studying the effect of heat (Figure 4 A and B), five TFFs (LSD, M-type MADS, RAV, SAP, and VOZ) were represented in analysis with microbes that were not present in the autoclaved, and three TFFs (HB-PHD, S1Fa-like, STAT), were unique to the autoclaved soil analysis. There were 176 differentially expressed TFs shared across the analyses. However, the majority of these were expressed in the opposite direction between the different soil microbiomes, with 74% being regulated in opposite directions. Only eight TFs were differentially expressed across all three genotypes in both soil microbiome analyses, with six of these shared TFs being regulated in opposite directions depending on soil type. This lends support that the interaction with the microbiome affects the response to heat stress in soybean. The two genes with the same expression profiles across microbiomes, *Glyma.15G029500* and *Glyma.17G227600*, encode for a DBB and HSF TF, respectively. The gene *Glyma.17G227600* was one of the few genes that was detected across all genotypes and soil type analyses, indicating its importance in response to heat in soybean. The *Arabidopsis* homolog (AT2G26150) is a member of the heat shock TFF (HsfA2) and increased expression of this gene confers thermotolerance, possibly through regulating JA synthesis (Guo et al., 2024). The gene *Glyma.15G029500* was downregulated across all genotypes and soil types. The *Arabidopsis* homolog of this gene (AT1G06040) is known to be important for conferring tolerance to high salt conditions (Chiriotto et al., 2023). It also plays an important role in light signaling as its expression is regulated by photoperiod.

#### 3.1.4. Characterization of Gene Co-expression Networks

We clustered similarly expressed genes that were differentially expressed for the three genotypes when compared to Williams82 into modules using weighted gene co-expression network analysis. For the effect of temperature, we created two co-expression networks based on the type of soil.

GO enrichment analysis was performed for each co-expression module gene list to provide biological insight into each module. Not all genes were assigned to a module in cases of low correlation for some genes. Hub genes for each module were determined as the genes that were the most highly connected.

The co-expression network for the effect of heat stress when grown in native microbial soil was built on 4224 DEGs. These genes were assigned to three separate modules, with 142 genes unassigned.

The largest module, with 2870 genes, contained genes associated with photosynthesis and plant immunity, as reflected by the 47 overrepresented GO terms. The second module had 938 genes and eight GO terms, most associated with the central dogma theory. The final module had 274 genes with 13 GO terms. The terms in this module include response to heat (GO:0009408) and other terms related to protein folding and import. This module also had *Glyma.17G227600* as one of the hub genes.

The second co-expression network was for the effect of heat stress when plants were grown in autoclaved soil and was built on 964 genes. These genes were also assigned to three modules, with 63 genes unassigned. The first module had 426 genes and had five overrepresented GO terms, which was dominated by terms related to photosynthesis. The second module had 393 genes, and 24 GO terms related to photosynthesis, and responses to different lights and heat. The HSF gene, *Glyma.17G227600*, was the top hub gene for this module. The final module for this network had 82 genes and only two GO terms (GO:0019375, galactolipid biosynthetic process; and GO:0016036, cellular response to phosphate starvation).

### 3.2. Effect of Soil Microbiome

#### 3.2.1. Identification and Comparison of DEGs across Genotypes

In addition to studying the effect of heat stress, the effect of the soil microbiome was studied for each genotype by comparing gene expression differences between plants grown in soil with a microbiome and plants grown in autoclaved soil (Figure 6). Similar to the heat stress response portion of the study, we conducted a separate analysis for each temperature treatment (heat vs control), as well as an analysis across the two temperature treatments. In these analyses, DEGs were only identified in a single genotype, PI89008. Under high heat conditions, we identified 6495 DEGs in PI89008 when comparing the effect of the microbiome in heat conditions, with 2794 of these genes upregulated in the absence of a microbiome and the remaining 3701 downregulated. Genes annotated as protein kinases made up 4.7% (304) of the DEGs in the heat analysis. The protein kinase genes were split, with 170 being upregulated in the absence of a microbiome and the remaining 134 being downregulated. A study in rice roots found that protein kinase domains were one of the core transcriptional responses due to changes in the soil microbiome (Santos-Medellín et al., 2022). 15 of the DEGs were also present in the combined temperature analysis, with nine of them being downregulated. No GO terms were significantly overrepresented among the 15 shared genes. Two of the 15 DEGs, *Glyma.08G009100* and *Glyma.18G287200*, encode for transcription factors from the bHLH and C2H2 TFFs, respectively. No DEGs were detected for microbiome effect when plants were grown in the optimal temperature. The response to microbiome under heat stress temperatures identified 59 significantly enriched overrepresented GO terms. Terms were largely described as processes related to DNA replication and cell multiplication and terms related to gene silencing. One GO term related to stress (response to salt stress (GO:0009651) was also significant.

**Figure 6.**
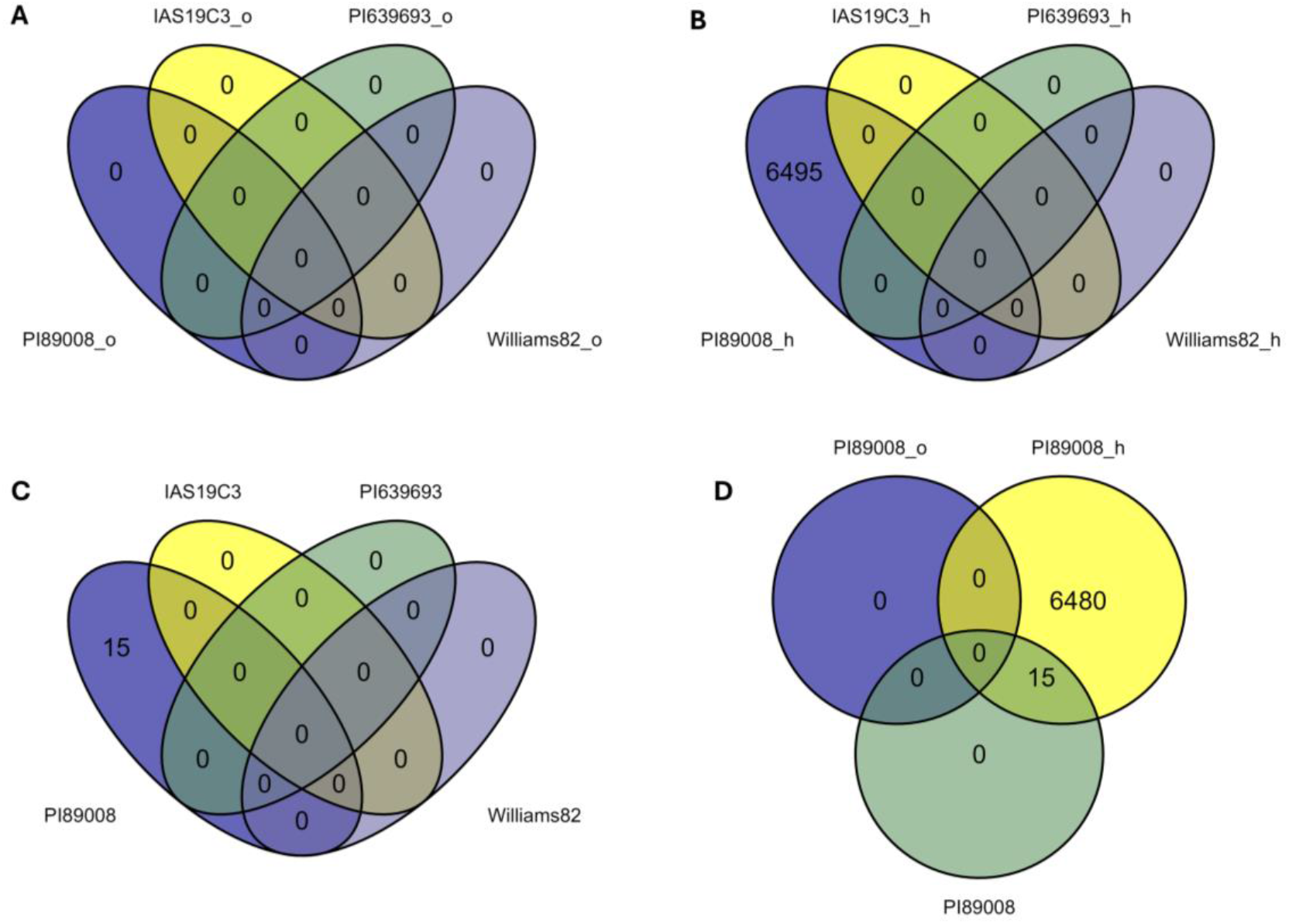
Venn diagrams of the distribution of overlapping significant DEGs (FDR < 0.05) due to the effect of a microbiome for each genotype. Here, we show that three genotypes had no significant DEGs detected for any of the temperature treatments, with (A) being the optimal temperature, (B) the heat stress temperature, and (C) the combined analysis across temperatures. Due to the lack of DEGs for three genotypes, Venn diagrams comparing DEGs between temperature analyses are not presented. The overlap of significant DEGs for PI89008 between temperature analyses is shown in (D).

#### 3.2.2. Identification and Comparison of DEGs between Genotypes

We compared the changes in the transcriptome between genotypes due to the effect of the microbiome, with three genotypes compared to Williams82 (Equations 3 & 4). In contrast to the comparisons within a single genotype (Figure 6), significant DEGs were detected for all genotypes in both temperature treatments (Figure 7). The overlap of DEGs between IAS19C3 and PI89008 was high, while the overlap between the three genotypes was lower, with 124 and 31 DEGs (Figure 7) in the heat stress and optimal temperature, respectively. Across both temperatures, DEGs shared between all genotypes were regulated in the same direction, with most being up regulated due to the presence of a soil microbiome. Additionally, we observed the DEGs from a single genotype that were shared between the temperature conditions. IAS19C3 had 78 DEGs differentially expressed under both temperature conditions, 72 of which are regulated in opposite directions. PI89008 had 401 DEGs differentially expressed under both temperatures, 399 of which are regulated in opposite directions. No DEGs were identified in PI639693 in both temperatures.

**Figure 7.**
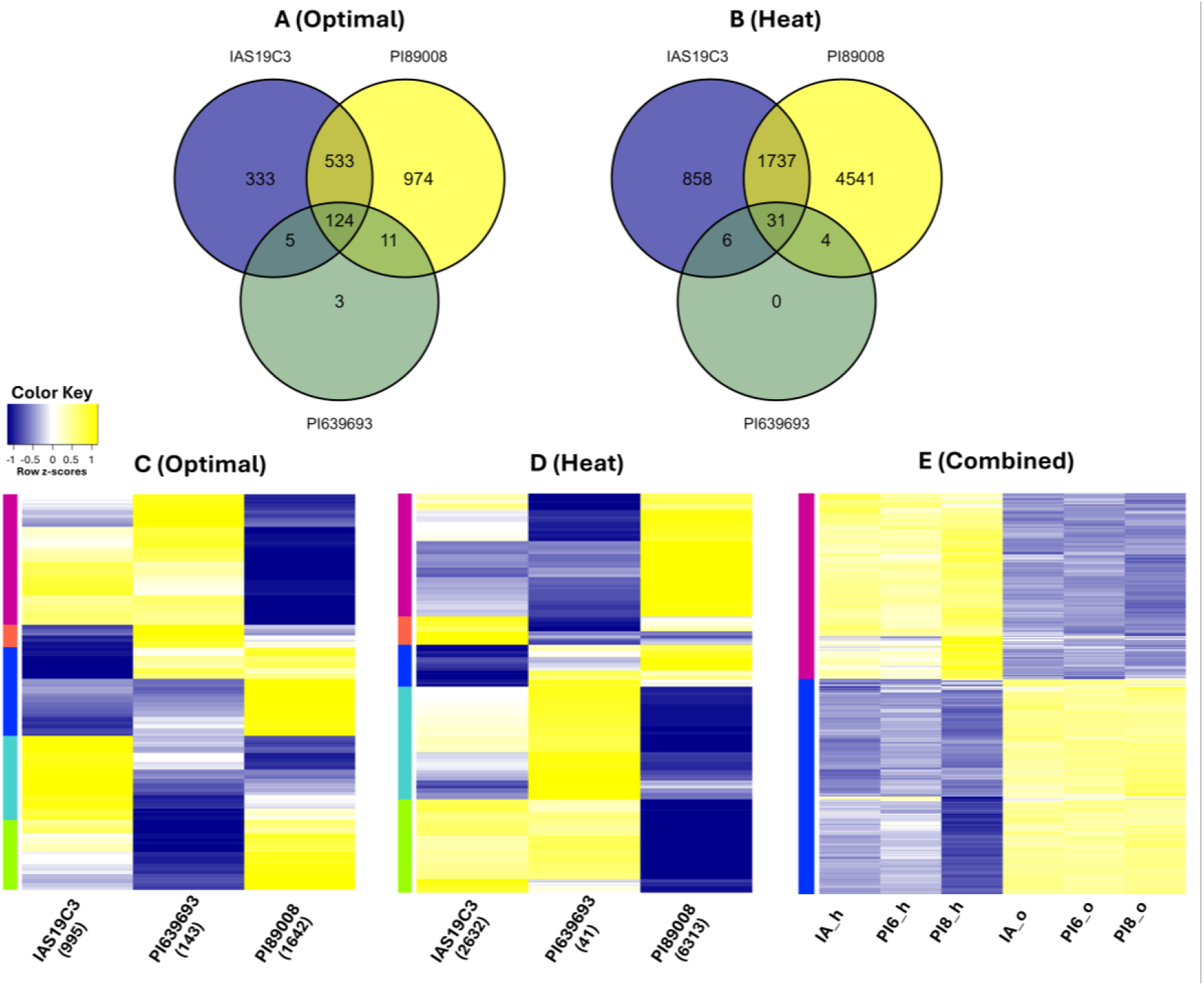
Distribution of differentially expressed genes (DEGs) and heatmaps of the DEGs due to the effect of a microbiome. The Venn diagrams are the distribution of statistically significant (FDR < 0.05) DEGs in each of the analyses, with (A) being the effect of microbiome and genotype when plants were grown in optimal temperatures and (B) being when the plants were grown in heat stress temperatures. Genotypes were compared to the control of Williams82 in autoclaved soil (no microbiome). The heatmaps demonstrate the expression levels of DEGs across all genotypes and analyses, with (C) being the DEGs of the effect of microbiome and genotypes with plants grown in optimal temperature, (D) when the plants were grown in heat stress temperatures, and (E) the DEGs across the optimal and heat soil analyses. Expression of all DEGs identified for each analysis is represented, even if a gene was not statistically differentially expressed in a specific comparison. The number of significant DEGs for a single genotype are in parentheses below the genotype name. Downregulated genes (negative Z-scores) are shown in blue, and upregulated genes are shown in yellow. Clusters of genes with similar expression profiles are outlined in colors to the left of each heatmap.

Again, we focused our analyses on the DEGs significant for all three genotypes in a single temperature. Of the 124 DEGs shared by all three genotypes under optimal temperature, nine DEGs were related to phosphate uptake and accumulation. However, this could be a result of increased available phosphorous in soil after autoclaving. Another fifteen genes are related to circadian rhythms,

including *Glyma.13G199300*, which is homologous with the *Arabidopsis* gene LNK2, which is involved in a regulatory mechanism for circadian signaling pathways (Sorkin et al., 2023). Several upregulated DEGs were homologous to *Arabidopsis* genes related to defense responses and immunity to pathogens, as well as to abscisic acid (ABA) production. The enrichment of these 124 genes resulted in eight significantly overrepresented GO terms. These terms were related to circadian rhythms, phosphates, and lipid biosynthetic processes. In contrast, fewer DEGs were shared between all three genotypes when grown in the heat stress conditions, with enrichment of the 31 genes resulting in ten overrepresented terms related to growth and development. Most of the 31 DEGs were homologous to *Arabidopsis* genes related to meristem formation, organ separation, and flowering. One gene of note, *Glyma.19G014400*, was homologous with *At5-MMP*, which is involved in recruiting differential root bacteria communities in

*Arabidopsis* (Mishra et al., 2021). This gene was downregulated in the presence of a soil microbiome, with a log fold change of 12-19, depending on the genotype in leaves, suggesting either an alternative function or leaves are important for initiating root signals. This gene could also be downregulated in the presence of a microbiome due to sufficient microbial communities already existing around the roots.

#### 3.2.3. Differentially Expressed Transcription Factors due to Microbiome Presence

Differentially expressed transcription factors were flagged for further analysis. Similar to the TF analysis for the effect of temperature, the log-fold change of the differentially expressed TFs were plotted by grouping the TFFs by genotype and temperature treatment, as shown in Figure 8. Under optimal temperatures, there were 210 differentially expressed TFs belonging to 37 unique TFFs. The majority of these TFs (69%) were differentially expressed in only a single genotype, primarily in PI89008. Only 6% of the TFs were differently expressed in all three genotypes, with all 13 TFs detected in PI639693 found in the other two genotypes as well. The presence of a microbiome resulted in the majority (74%) of the TFs in optimal temperatures being downregulated. More differentially expressed TFs were detected under heat stress conditions, with a total of 579 TFs from 48 TFFs. Trends in expression were similar to the optimal conditions, with the majority of the TFs (69%) being differentially expressed in only a single genotype. PI639693 had only 16 differentially expressed TFs, and 14 of these were found to be differentially expressed for all three genotypes. In contrast to the optimal

**Figure 8.**
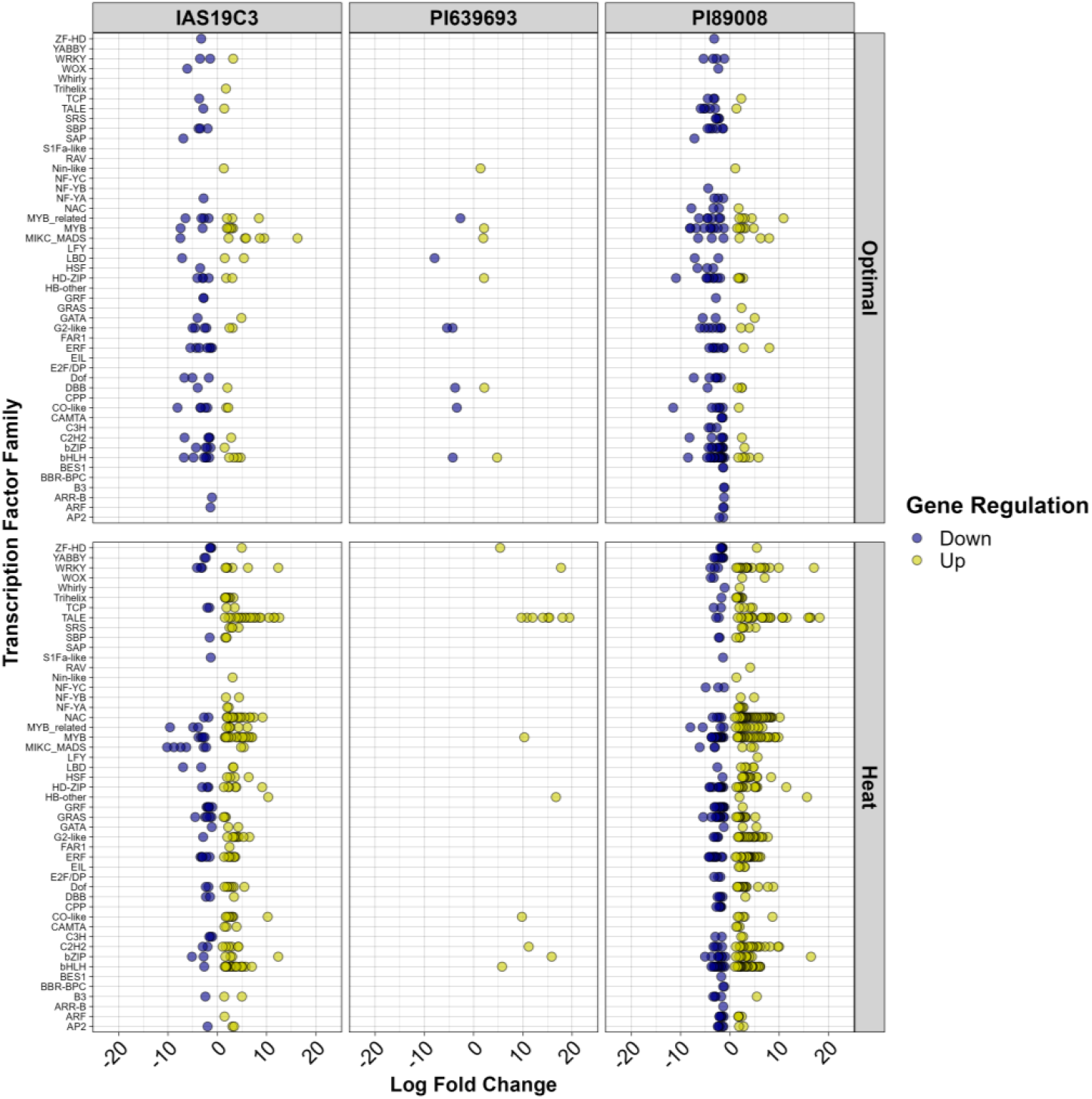
Expression values of differentially expressed transcription factors of three soybean genotypes compared to the control genotype Williams82. Differentially expressed genes were identified in the leaf tissue of plants in response to differences in the soil microbiome. Plants grown in the differing microbiomes were also grown in two different temperature conditions, optimal and heat stress. Transcription factors were identified in each DEG list and plotted by transcription factor family using the log fold-change of the gene. Up-regulated genes are shown in yellow, while down-regulated genes are shown in blue.

temperatures, in heat stress, the presence of a microbiome resulted in 387 TFs (67%) being upregulated.

A single TFF, SAP, was found to be differentially expressed in optimal temperatures (in IAS19C3 and PI89008) and not in the heat stress conditions. An additional 12 TFFs were only differentially expressed due to the microbiome effect when grown in heat stress conditions. Between the two temperatures, there were 62 shared differentially expressed TFs. Interestingly, all TFs were regulated in the opposite direction when compared across temperatures, with the majority (71%) being downregulated due to the presence of a microbiome in the optimal temperature and then upregulated when in the heat stress conditions.

## 4. Discussion

Heat stress is already a major abiotic stress affecting yield in numerous field crops and is predicted to become more important in soybean by the end of the century (Yang and Wang, 2023). To combat the effects of heat stress, it is important to identify tolerance mechanisms in crops.

Transcriptomic analysis of plant responses to different stresses is one effective approach to identifying key tolerance genes and pathways for stress response. Additionally, a growing body of evidence indicates that soil microorganisms may benefit plants that are experiencing abiotic stress and help alleviate the symptoms (Caddell et al., 2019). The effect of short-term heat stress in soybean has been previously investigated via transcriptomics (Song et al., 2017; Valdés-López et al., 2016; Wang et al., 2018). To our knowledge, no report has focused on the transcriptome of continuous and long-term heat stress in soybean. Additionally, few studies have reported on the effect of a microbiome in crop species (Santos-Medellín et al., 2022), especially in combination with abiotic stress conditions (Timm et al., 2018).

Therefore, in this study, we focused on changes in the transcriptome of soybean due to heat stress and due to a microbiome and how these changes affected each other. This will help identify novel stress tolerance mechanisms to exploit in soybean breeding and genetics to develop more climate resilient cultivars.

We aimed to identify DEGs under heat stress in different soil microbiome conditions. Across the three genotypes that were compared to Williams82, several thousand unique genes were identified.

Therefore, we found it helpful to focus on the DEGs that were shared between all three of the genotypes. We hypothesized the overlapping DEGs may be major regulators of soybean response to heat. In total, we identified 4224 overlapping DEGs due to heat stress in native soils and 964 overlapping DEGs when there was no soil microbiome. These overlaps accounted for 40.8% and 9.4% of the total DEGs detected in response to heat for both the presence and absence of a microbiome, respectively. The low proportion of overlapping DEGs between genotypes is consistent with previously reported results in *Arabidopsis* (Barah et al., 2013) and wheat (Rangan et al., 2020) exposed to heat stress, as well as soybean to iron deficiency stress (Kohlhase et al., 2021), indicating the differential responses to stress that unique genotypes may have and supporting the need to study multiple genotypes. Additionally, there were only 83 DEGs that were shared between all three genotypes and across the soil microbiome treatments, which indicates that soil microbiomes are contributing to soybean response to heat stress. The 83(?) shared genes were largely related to both abiotic and biotic stress responses, including the synthesis of ABA and numerous HSPs. In responses to heat in two different soil microbiome conditions identified genes related to photosynthesis were more prominently downregulated when there was no soil microbiome, as opposed to when there was a soil microbiome. However, the gene networks constructed for both soil microbiome types had modules with genes largely associated with photosynthesis, indicating that photosynthetic processes are greatly affected by heat stress.

A promising gene, *Glyma.17G227600*, was upregulated in all three genotypes due to heat, regardless of genotype or soil microbiome. This gene is an HsfA2 and was also found to be a hub gene with many co-expression correlations to other genes during the network analysis for the effect of temperature. The HSF family has been important in researching heat stress tolerance in many plant species (Fragkostefanakis et al., 2015; Ohama et al., 2017). Specifically, HsfA2s are essential for the heat stress response in plants and are hypothesized to be involved with the acquisition of thermotolerance by regulating the expression of chaperone proteins and involved in heat shock memory for a long-term adaptation to heat stress (Ohama et al., 2017). In soybean there are five genes that encode for HsfA2 (Sinha et al., 2023). Given that the gene *Glyma.17G227600* was found to be upregulated across genotypes and soil microbiome conditions, we hypothesize this gene to be critical for soybean heat stress tolerance.

An additional 24 HSFs, including three HsfA2s, were differentially expressed in at least one comparison, but no other HSFs were conserved across all genotypes and soil conditions. The gene *Glyma.17G053700*, another HsfA2, was the next most conserved HSF, was downregulated in all genotypes when grown in microbial soil, and upregulated for one genotype in the autoclaved soil. The remaining two HsfA2s, *Glyma.04G052000* and *Glyma.14G096800* were upregulated in response to heat for all genotypes when grown in autoclaved soil, but were not differentially expressed when in microbial soil. The majority of HSFs were upregulated due to heat stress when the plants were grown with no microbiome, while they were downregulated when grown with a microbiome (Figure 5). This could be due to beneficial microorganisms alleviating some of the symptoms of heat stress, negating the need to activate HSFs. A previous study found that inoculation of soybean exposed to 5-10 days of heat stress with a specific *Bacillus* strain resulted in protection of photosynthetic pigments and augmented heat stress responses (Khan et al., 2020).

We characterized the changes in the soybean transcriptome comparing whether a microbiome was present under both optimal and heat stress conditions. Phenotypic measurements show plants grown with no soil microbiome mature later, indicating the assistance of a soil microbiome in early plant growth and development. Comparing analyses from the different temperature treatments, no significant DEGs were shared across all genotypes. Different temperatures are likely to promote the growth of variable microbial communities, and the lack of shared genes could be due to the different communities present across temperatures. Within a single temperature condition, few genes were shared across the genotypes, indicating the possibility of genotypic specific responses to the microbiome. A common set of genes in optimal temperatures were related to phosphorus deficiency and accumulation. We credit this in part due to the higher amounts of available phosphorus in the autoclaved soil, which was hypothesized to be due to the breakdown of organic matter due to the temperature and pressure when autoclaving (personal communications). Other core genes between the genotypes were largely related to circadian clocks and plant immunity. The soil microbiome is likely to contain both pathogenic microbes and beneficial microbes, although we noted no visible diseases on our plants. Plant immunity to microbes is complex, and it has been proposed that the response is not only against pathogens but also being used to establish and maintain homeostasis with the root microbiome (Thoms et al., 2021).

The effect of the soil microbiome, when plants were grown in heat, had even fewer DEGs commonly shared across all genotypes, with only 31 shared genes. In contrast to the optimal temperature analysis, none of the shared genes were related to phosphorus, even though the soils had the same levels of phosphorus at the time of planting as the optimal temperature condition. We speculate that this could be due to the prevalent microbes in the heat stress conditions using less phosphorus, thus less competition with the plants. The core set of genes found in this analysis was largely related to meristem and organ development and maintenance. This supports the hypothesis that the soil microbiome plays a role in promoting growth and development in plants.

## 5. Conclusions

Our work shows that many genes are differentially expressed in response to long-term heat stress and that different genotypes have unique responses. However, there are conserved DEGs that may be common across genotypes. The results also demonstrate that the presence or absence of a soil microbiome does affect the transcriptional response to heat stress in a significant way. The limited number of DEGs shared between genotypes and soil microbiome status may play key roles in the physiological function of soybean response to long-term heat stress. Additionally, we have shown that soybean transcriptomes respond to the presence of a soil microbiome and that the response to the microbiome is unique between soybean genotypes. This response to the microbiome is also affected by the temperature of the environment, and this microbiome may assist in alleviating some symptoms of abiotic stress. To our knowledge, this is the first study in soybean on changes in the transcriptome under long-term heat stress, as well as the first observing changes due to the microbiome. This research helps identify genes for plant stress tolerance, and in determining how the interaction with the microbiome may play a role in the stress responses, thus lending future directions for breeding of heat tolerant soybean.

## Acknowledgments

We thank members of the AKS group at ISU for their technical support. We thank Drs. Rex Nelson and Jaqueline Campbell for assistance with SoyBase. We thank Aaron Brand for his assistance in preparing the soil. We thank Scott Zarecor and Dr. Steve Whitam for the scheduling and programming of the growth chambers in the Enviratron.

## Data Availability Statement

Data will be made available with publication of paper in peer reviewed journal.

## Author Contributions

LV: Conceptualization, Methodology, Software, Validation, Formal analysis, Investigation, Writing – original draft, Writing – review & editing, Visualization. DE: Methodology, Investigation, Writing - review & editing. AF: Methodology, Writing – review & editing. JAO: Methodology, Software, Validation, Formal analysis, Writing – review & editing, Visualization. AKS: Conceptualization, Methodology, Resources,

Writing – review & editing, Supervision, Project administration, Funding acquisition.

## Funding

The authors sincerely appreciate the funding support from the Iowa Soybean Research Center, Raymond F. Baker Center for Plant Breeding, Plant Sciences Institute, and the G.F. Sprague Chair in Agronomy. Additional funding from USDA-ARS project 5030-21220-007-000D.

**Table 1.** Average vegetative growth stages of each genotype in each factorial treatment.

| Genotype | Temperature | Soil Microbiome |  |
| --- | --- | --- | --- |
|  |  | Native | Autoclaved |
| Williams82 | OT | 6 | 4 |
|  | HT | 6 | 5 |
| IAS19C3 | OT | 6 | 4 |
|  | HT | 6 | 6 |
| PI639693 | OT | 7 | 5 |
|  | HT | 7 | 5 |
| PI89008 | OT | 7 | 5 |
|  | HT | 8 | 6 |

